# Characterization of mechanical tissue properties in post-mortem human brain using magnetic resonance elastography

**DOI:** 10.1101/2025.11.12.688069

**Authors:** Joy Mojumder, Yuan-Chiao Lu, Alexa M. Diano, Ahmed A. Alshareef, Matthew McGarry, Philip V Bayly, Curtis L. Johnson, John A. Butman, Dzung L. Pham

## Abstract

Traumatic brain injury (TBI) is a serious health condition that can cause neurological dysfunction to varying degrees depending on the nature of the mechanical insult. In biomechanical studies of TBI under high loading conditions, post-mortem human subjects (PMHS) are often used since ethical concerns prohibit such experiments in living human subjects. Because PMHS brains undergo significant changes following death, it is important to understand the relationship between the mechanical properties of PMHS brain tissue and living tissue. In this study, we performed magnetic resonance elastography (MRE) on three PMHS specimens to estimate the material properties of the cadaveric brain, namely the storage modulus and the loss modulus, as well as the resulting shear stiffness and damping ratio. We also performed longitudinal MRE scans on one of the PMHS over the span of two months to investigate the evolution of tissue properties with post-mortem degradation. In comparison to *in vivo* subjects of age range 70-75 years, a substantially higher stiffness (mean: 5.96kPa) and lower damping ratio (mean: 0.09) were found in PMHS models. This study also revealed an initial increase in shear stiffness up to the seventh day post-mortem, followed by a steady decrease by the fifty-eighth day. However, the damping ratio displayed an opposite trend to that of shear stiffness. These changes were heterogeneous across brain regions. The collected measurements and analysis elucidate the changes in mechanical properties in post-mortem subjects, and can be used to build and validate computational models of TBI.

## 1. Introduction

The brain is a complex and important organ that undergoes anatomical and physiological changes from infancy to old age (Budday, Raybaud, et al., 2014; Budday, Steinmann, et al., 2014), and may also incur pathological alterations due to disease (Tyler, 2012). Traumatic brain injury (TBI) is a serious health condition caused by external mechanical forces, affecting tens of millions across both military and civilian populations, and is most prevalent in people younger than 25 and older than 75 years (Peeters et al., 2015; Silverberg et al., 2020). Based on the nature of the mechanical insult, the severity of TBI can be classified as mild, moderate, or severe in outcome. An estimated 42 million people sustain mild TBI (mTBI) each year globally, the most common type of TBI (Gardner & Yaffe, 2015). TBI across the severity spectrum can lead to debilitating persistent symptoms, including headaches, sleep dysfunction, and cognitive impairment. TBI patients are also more likely to develop neurodegenerative diseases, such as Alzheimer’s disease, Parkinson’s disease, and amyotrophic lateral sclerosis (Gardner & Yaffe, 2015).

Because treatments for TBI remain limited(Min & Shin, 2022), prevention is an important approach to limiting its negative consequences. Protective equipment and safety standards rely on computational models of TBI that simulate the biomechanical response of the brain under a range of loading conditions (Rowson & Duma, 2022). Several computational models have been developed, providing insights into both normal and diseased brain mechanics (Cloots et al., 2011; Feng et al., 2016; Giordano & Kleiven, 2014; Giudice et al., 2019; Kyriacou et al., 2002; Wright et al., 2013). While these computational models are a great tool for investigating the intrinsic mechanics of brain tissue, the implementation of these models depends on multiple factors, including the geometry, boundary conditions at tissue interfaces, and material properties. Furthermore, validation of these models requires experimental displacement and/or strain data, which is very scarce. Thus, the role of experimental studies in informing the mechanical properties of brain tissue in computational models becomes paramount.

Experimental studies have been performed in animal models and in humans *in vivo* to investigate the mechanical behavior of the tissues (Budday et al., 2017; Gefen & Margulies, 2004; Miller et al., 2000; Wang et al., 2024; Whittall et al., 1997). However, animal models do not have the same anatomical geometry, while human *in vivo* studies are limited to non-injurious loading conditions. In addition, *in vitro* and *in situ* post-mortem studies have been performed. For example, one study using indentation testing found *in vivo* brain stiffness to be on the same order of magnitude as *in vitro* measurements in porcine model (Miller et al., 2000; Wang et al., 2024), whereas another study reported stiffer brain tissue *in vivo* compared to *in situ* tests performed on rat model (Gefen & Margulies, 2004). While these studies offer insights into post-mortem changes in brain tissue, the applicability of the results to the human brain is limited due to the anatomical differences between animals and humans, the constraints of the indentation method, and the disturbance of the brain’s physical and chemical environment caused by skull removal and brain exposure. To overcome these limitations, post-mortem human specimens (PMHS) have been widely used to elucidate the biomechanical response of the brain at injury-level loading (Hardy et al., 2007; Singh et al., 2024). However, the relationship between the mechanical properties of a living human brain and that of a cadaver remains poorly understood.

Several non-invasive techniques, e.g., tagged magnetic resonance (MR) imaging and MR elastography (MRE) (Arani et al., 2021; Di Ieva et al., 2010; Murphy et al., 2019; Yin et al., 2018), have been developed to investigate the *in vivo* mechanical response of the human brain. MRE enables estimates of the material properties of the brain by applying harmonic loading to the skull during scanning, measuring brain tissue displacement, and then solving an inverse problem to estimate the storage and loss moduli (Bayly et al., 2021; Di Ieva et al., 2010). MRE has also been performed on animal models and phantom models to investigate the mechanical properties of brain tissue, as well as brain mechanics. For example, results from MRE performed on minipigs and rats showed that the stiffness of brain tissue increased directly following death (Gefen & Margulies, 2004; Miller et al., 2000). However, there is a lack of MRE data on the post-mortem human brain. Moreover, no study has investigated how the properties of post-mortem human brain tissue change *in situ* and over an extended period after death.

To investigate the mechanical properties of brain tissue in PMHS, we conducted MRE on three PMHS heads within 72 hours after death. Additionally, we performed longitudinal MRE scans on one of the specimens up to 58 days after death to examine changes in tissue properties over time. Shear waves were induced along the anterior-posterior (AP) direction of the PMHS models. Using the nonlinear inversion (NLI) technique (McGarry et al., 2022), we estimated various material properties of brain tissue both globally and regionally and compared them to *in vivo* measurements, elucidating the evolution of brain tissue material properties post-mortem.

## 2. Materials and methods

### 2.1 Post-mortem human specimens (PMHS)

Three PMHS heads were obtained through the Maryland State Anatomy Board (Baltimore, Maryland, USA). All PMHS information were summarized in **Table 1**. The neck was severed and sealed at approximately the fourth cervical vertebra to preserve fluid in the intracranial cavity. PMHS 1 was scanned at Days 2, 3, 5, 7, 9, 16, 20, 30 and 58 after death to investigate alterations in the mechanical properties over time. The heads were stored at 4^°^C in between scanning sessions to limit tissue degradation. Approximately 2-3 hours before scanning, the PMHS heads were removed from refrigeration to allow them to reach room temperature.

**Table 1.** PMHS information.

| Index | PMHS 1 | PMHS 2 | PMHS 3 |
| --- | --- | --- | --- |
| Sex | Female | Male | Female |
| Age (Years) | 91 | 91 | 79 |
| Approximate scan interval from time of death (days) | 2, 3, 5, 7, 9, 16, 20, 30, 58 | 3 | 2 |

### 2.2 MRI Scans

MRI was performed with a 16-channel receive-only head/neck coil in the Clinical Center at the National Institutes of Health (Bethesda, Maryland, USA). T1-weighted (T1-w) scans used Magnetization Prepared Rapid Gradient Echo (MPRAGE) sequences and were acquired to define anatomical regions of interest. The T1-w scan for PMHS 1 was acquired on a Siemens Aera 1.5T (TR=2200ms, TE=2.65ms, TI=1100ms, FA=7⁰, voxel size=1×1×1mm, matrix size=256×256×192), while PMHS 2 and 3 were acquired on a Siemens Biograph mMR 3T (TR=2530ms, TE=3.03ms, TI=900ms, FA=8⁰, voxel size=1×1×1mm, matrix size=256×256×192).

During the MRE experiment, the heads were placed on a soft “pillow” driver attached to a pneumatic actuator system (Resoundant Inc., Rochester, MN, USA) that applied an excitation at 50 Hz in the anterior-posterior (AP) direction (shown in **Figure 1a-1c**). Using a single-shot echoplanar imaging (EPI) sequence covering the entire brain, MRE displacement data were acquired at 50 Hz frequency on a Siemens Aera 1.5T (TR=9607ms for PMHS 1 and 6720ms for PMHS 2 and 3, TE=113ms for PMHS 1 and 69ms for PMHS 2 and 3, pixel size=2.5×2.5mm, slice thickness=2.5mm).

**Figure 1.**
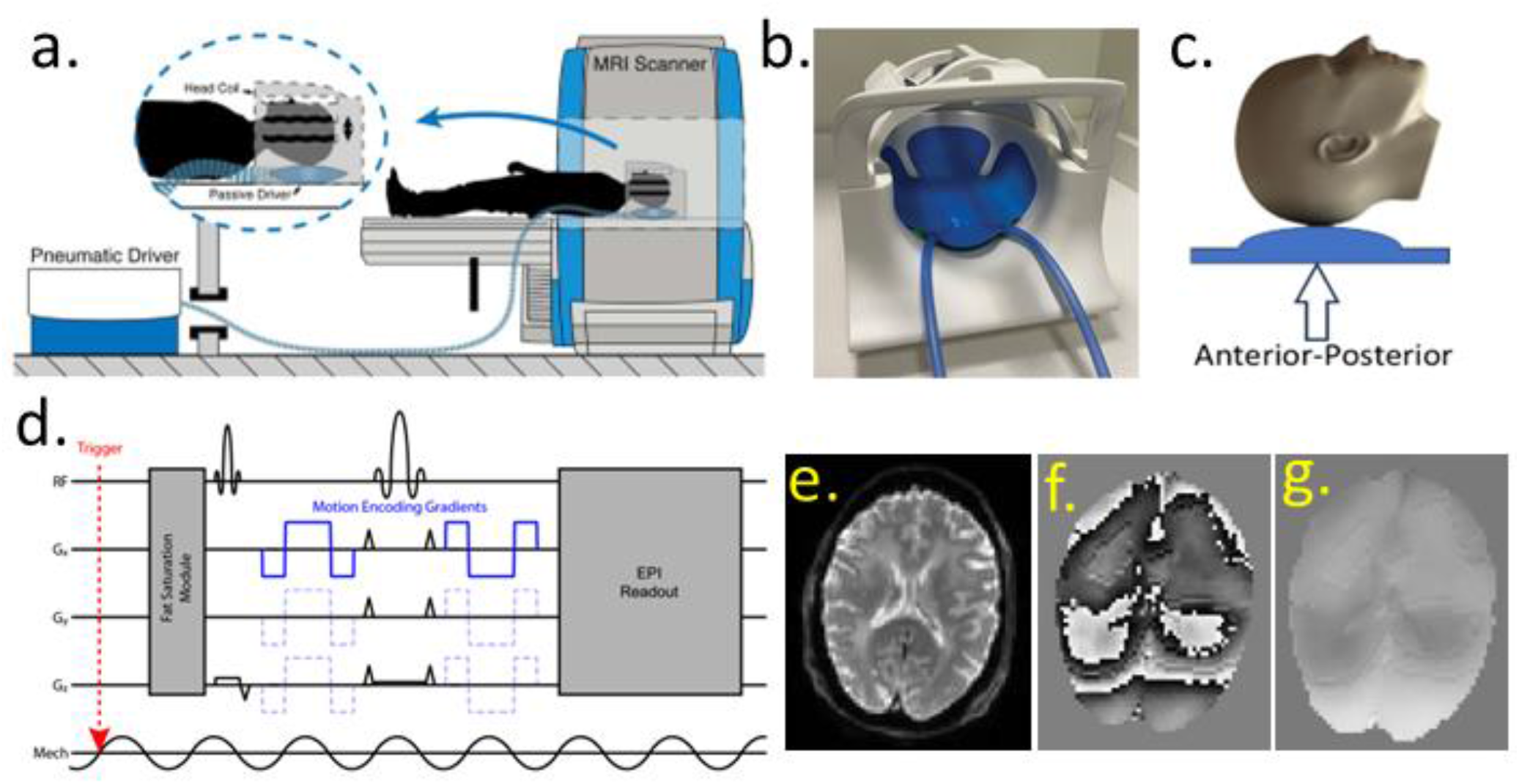
(a) Schematic of a typical experimental setup for MRE. (b) Pillow driver used for mechanical excitation, with induced motion along the anterior-posterior (AP) direction illustrated in (c). (d) Representative echoplanar imaging sequence incorporating motion encoding gradients. (e) Magnitude image, (f) wrapped phase image, and (g) unwrapped phase image acquired from the MRE experiment at 50 Hz in the central slice of the PMHS 1 model.

### 2.3 Data processing

Phase-contrast images encoding harmonic displacement were spatially unwrapped using PRELUDE (Smith et al., 2004) from the FMRIB Software Library (FSL) (Jenkinson, 2003) and were temporally Fourier transformed to obtain the frequency domain complex displacement field (Bayly et al., 2021; McGarry et al., 2022). The displacement data were used as input for the NLI algorithm (McGarry et al., 2022), a finite element-based optimization approach to estimate brain tissue material properties. NLI iteratively updates a set of mechanical property parameters from a heterogeneous finite element model to minimize the difference between model predictions and MRE tissue motion measurements. Considering brain tissue as heterogeneous, incompressible, and viscoelastic, the complex shear modulus, *G*^∗^ estimated by NLI is described as follows:

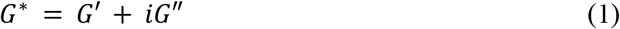

where *G*′ and *G*″ are the storage modulus and the loss modulus, respectively, representing the elastic and viscous behavior of tissue. Maps of *G*′ and *G*″ are recovered voxel-wise at the same resolution as the MRE data inputs. The complex modulus was converted to the shear stiffness *μ*, and damping ratio *ξ*, given by,

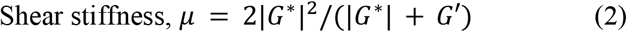

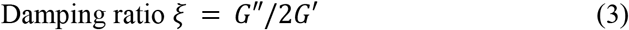

To obtain regional statistics of mechanical properties from NLI, the whole brain was segmented using a deep learning-based segmentation algorithm (Spatially Localized Atlas Network Tile – SLANT) from the T1-weighted images (Huo et al., 2019). After skull-stripping of T1-weighted images using mri_synthstrip (Hoopes et al., 2022), magnitude MRE images were affinely registered using the FLIRT command from FSL. Mechanical properties from NLI were then transformed to the T1 space, and regional masks were applied to quantify properties. SLANT labels were merged to form 5 major regions of interest: cerebellar white matter, cerebellar gray matter, cerebral white matter, cortical gray matter, and deep gray matter.

### 2.4 Statistical analysis

The mechanical properties of brain tissue in the PMHS models were reported by median ± median absolute deviation (MAD). The distribution of mechanical properties in PMHS models were compared to the properties estimated from *in vivo* subjects. Data of *in vivo* subjects within age range of 35 ± 15 years (32 subjects: Male = 14, Female = 18) and 73 ± 2 years (4 subjects : Male = 3, Female = 1) were collected from the brain biomechanics imaging resources (https://www.nitrc.org/projects/bbir) (Alshareef et al., 2025; Bayly et al., 2021). The distribution of mechanical properties in 5 different brain regions (cerebellar gray and white matter, cerebral gray and white matter, and deep gray matter) were reported after excluding the ventricles. Statistical significance of differences between groups was evaluated regionally using the Mann-Whitney U test.

## 3. Results

### 3.1 Whole brain mechanical properties in PMHS models and *in vivo* subjects

The T1 weighted anatomical images of the 3 PMHS are shown in Fig. 2a (Hoopes et al., 2022; Smith, 2002). The spatial distribution of normalized shear stiffness and damping ratio on an axial slice for all 3 PMHS is shown in Figures 2b and 2c. The median ±MAD of the mechanical parameters are reported in Table 2. The stiffness was highest and damping ratio was lowest in PMHS 2 (*μ* = 8.3 ± 1.80 *kPa, ξ* = 0.07 ± 0.02), which was scanned at a later time after death compared to PMHS 1 (*μ* = 4.8 ± 1.20*kPa, ξ* = 0.09 ± 0.03) and PMHS 3 (*μ* = 4.4 ± 0.80 *kPa, ξ* = 0.1 ± 0.04). Similar trends were also observed spatially in Figure 2b and 2c. Note that property estimates in the ventricles and all cerebrospinal fluid filled regions will not be accurate due to flow and were excluded in these measurements.

**Table 2.** Mechanical properties in PMHS models across the whole brain.

| Parameters | PMHS 1 | PMHS 2 | PMHS 3 | Age 20-50 | Age 70-75 |
| --- | --- | --- | --- | --- | --- |
| Shear stiffness (kPa) | $4.80 \pm 1.20$ | $8.30 \pm 1.80$ | $4.4 \pm 0.80$ | $2.80 \pm 0.40$ | $2.40 \pm 0.40$ |
| Damping ratio | $0.09 \pm 0.03$ | $0.07 \pm 0.02$ | $0.10 \pm 0.04$ | $0.20 \pm 0.10$ | $0.25 \pm 0.07$ |
| Storage modulus (kPa) | $4.60 \pm 1.20$ | $8.10 \pm 1.80$ | $4.2 \pm 0.80$ | $2.3 \pm 0.40$ | $1.90 \pm 0.45$ |
| Loss modulus (kPa) | $0.92 \pm 0.24$ | $1.27 \pm 0.36$ | $0.90 \pm 0.24$ | $1.10 \pm 0.24$ | $0.94 \pm 0.24$ |

**Figure 2.**
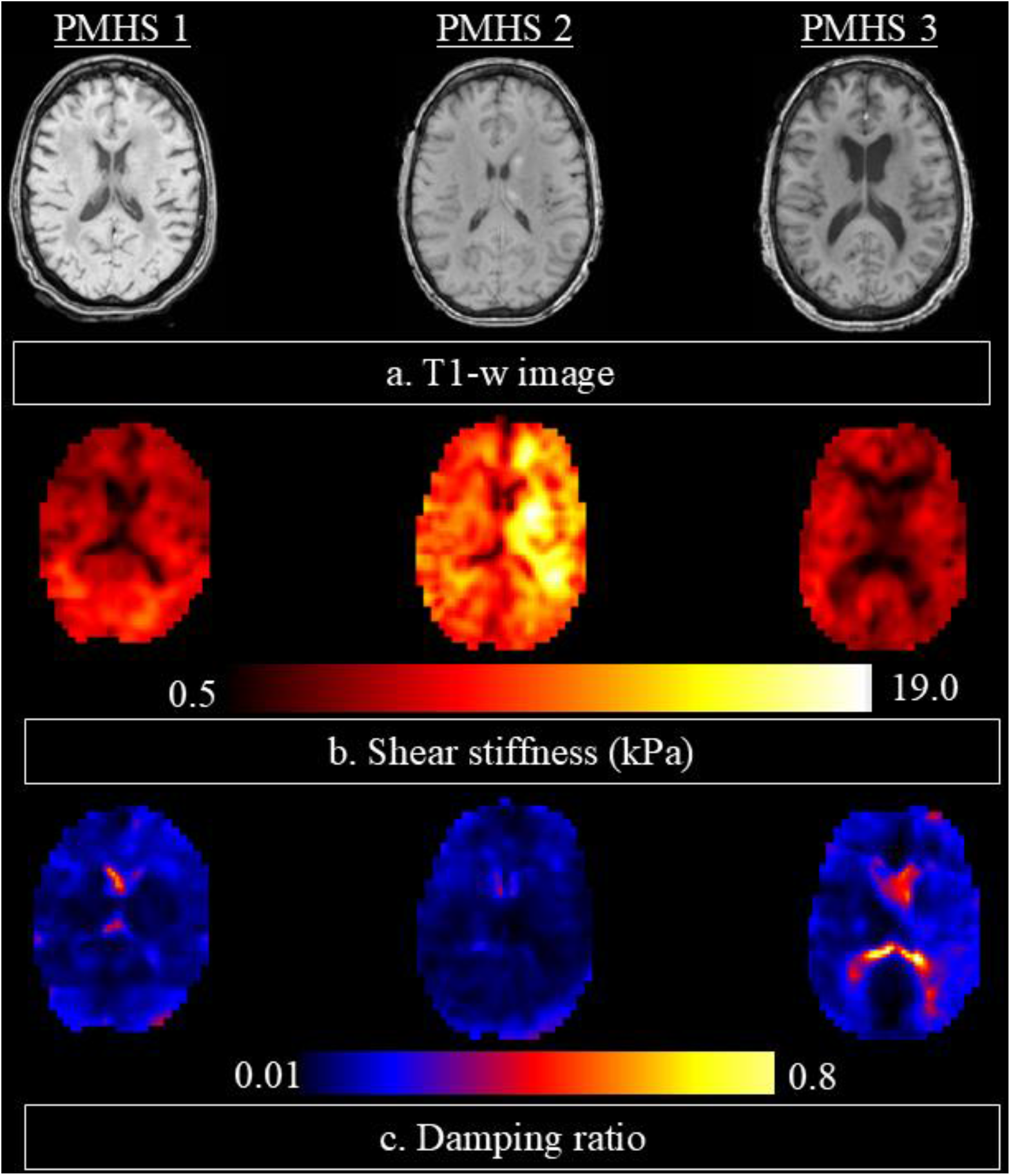
(a) Representative slices from T1-weighted scans for the PMHS 1 baseline, PMHS 2, and PMHS 3 scans. Qualitative comparisons of (b) shear stiffness and (c) damping ratio across the three PMHS models are presented.

The distributions of estimated mechanical properties across the whole brain for the 3 PMHS were compared to the corresponding properties of *in vivo* subjects (Fig. 3a – 3d). Quantitatively, notably higher stiffness was found in PMHS 2 (∼245%), followed by PMHS 1 (∼104%), when compared to the *in vivo* subjects with age range 70-75 years. However, a notably lower damping ratio was found in PMHS 2 (∼73% lower) followed by PMHS 1 (∼61% lower) and PMHS 3 (∼56% lower) compared to the *in vivo* subjects.

**Figure 3.**
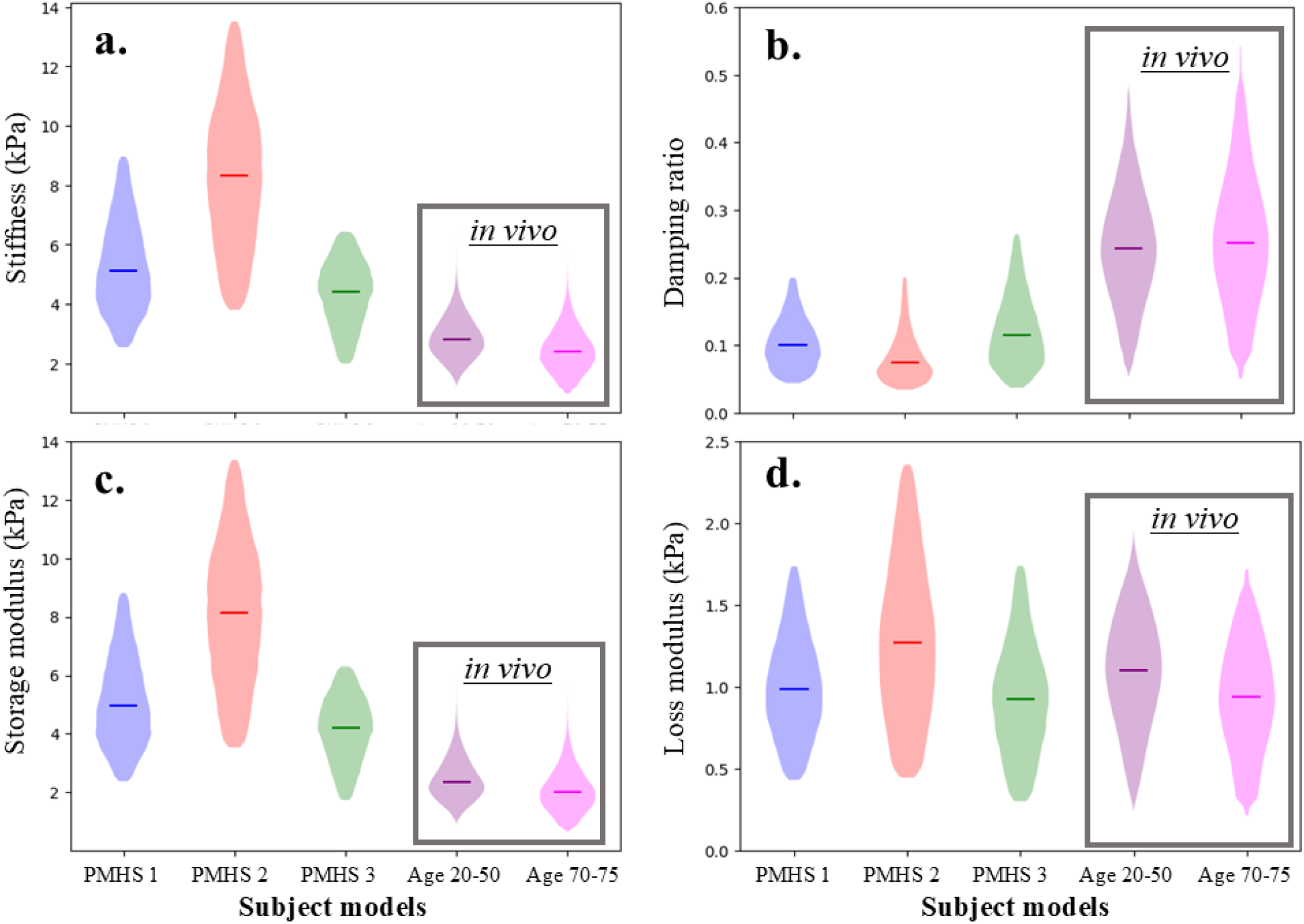
Violin plots of (a) stiffness, (b) damping ratio, (c) storage modulus, and (d) loss modulus of the whole brain for three PMHS models, compared to in vivo data from human subjects aged 20-50 years and 70-75 years. In vivo data are presented in black boxes.

Figure 4 compares stiffness and damping ratio of the PMHS and *in vivo* subjects across white and gray matter regions of the brain. The stiffness of white matter (WM) was slightly higher than that of gray matter (GM) in PMHS 1 (WM : 5.3 ± 1.01 *kPa* vs GM: 4.8 ± 1.16 *kPa*) and PMHS 2 (WM: 9.3 ± 1.53 *kPa* vs GM: 7.6 ± 1.66 *kPa*), whereas white matter was found to be softer compared to gray matter in the PMHS 3 (WM: 4.5 ± 0.7 *kPa* vs GM: 4.56 ± 0.8 *kPa*) model (Figure 4a). On the other hand, the damping ratio was slightly higher in white matter compared to gray matter in PMHS 1 and PMHS 3 model, whereas damping ratio was slightly lower in white matter compared to gray matter in PMHS 2 model (Fig. 4).

**Figure 4.**
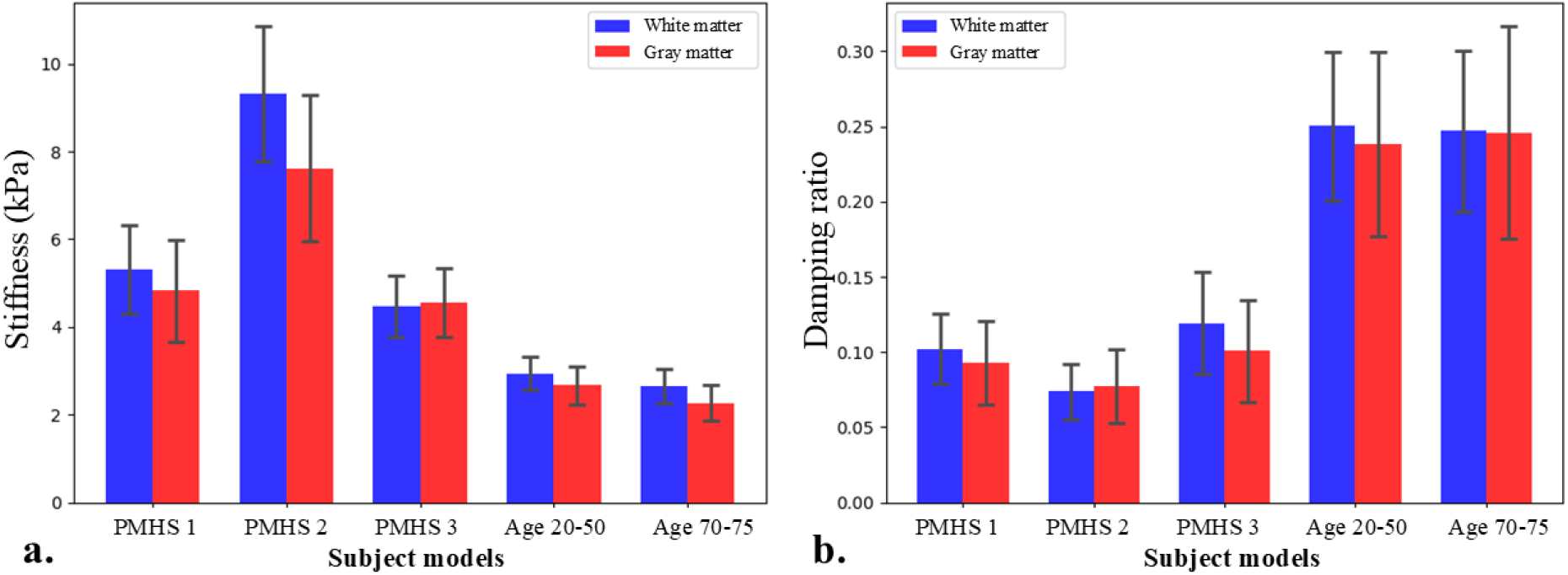
Comparison of (**a**) stiffness and (**b**) damping ratio between white and gray matter across three PMHS models and *in vivo* subjects.

### 3.2 Regional analysis of mechanical properties

Violin plots of mechanical properties of more specific brain regions are shown in Fig. 5. The quantitative values of these mechanical properties are reported in Table A.1 and A.2. Cerebral regions (cerebral white matter, cortical gray matter, deep gray matter) showed similar patterns to the whole brain results, with PMHS 2 exhibiting higher stiffness and lower damping ratios than PMHS 1 and 3. However, in cerebellar regions the patterns were different. PMHS 1 had the highest stiffness and lowest damping ratio in the cerebellum, while PMHS 3 had the lowest stiffness.

**Figure 5.**
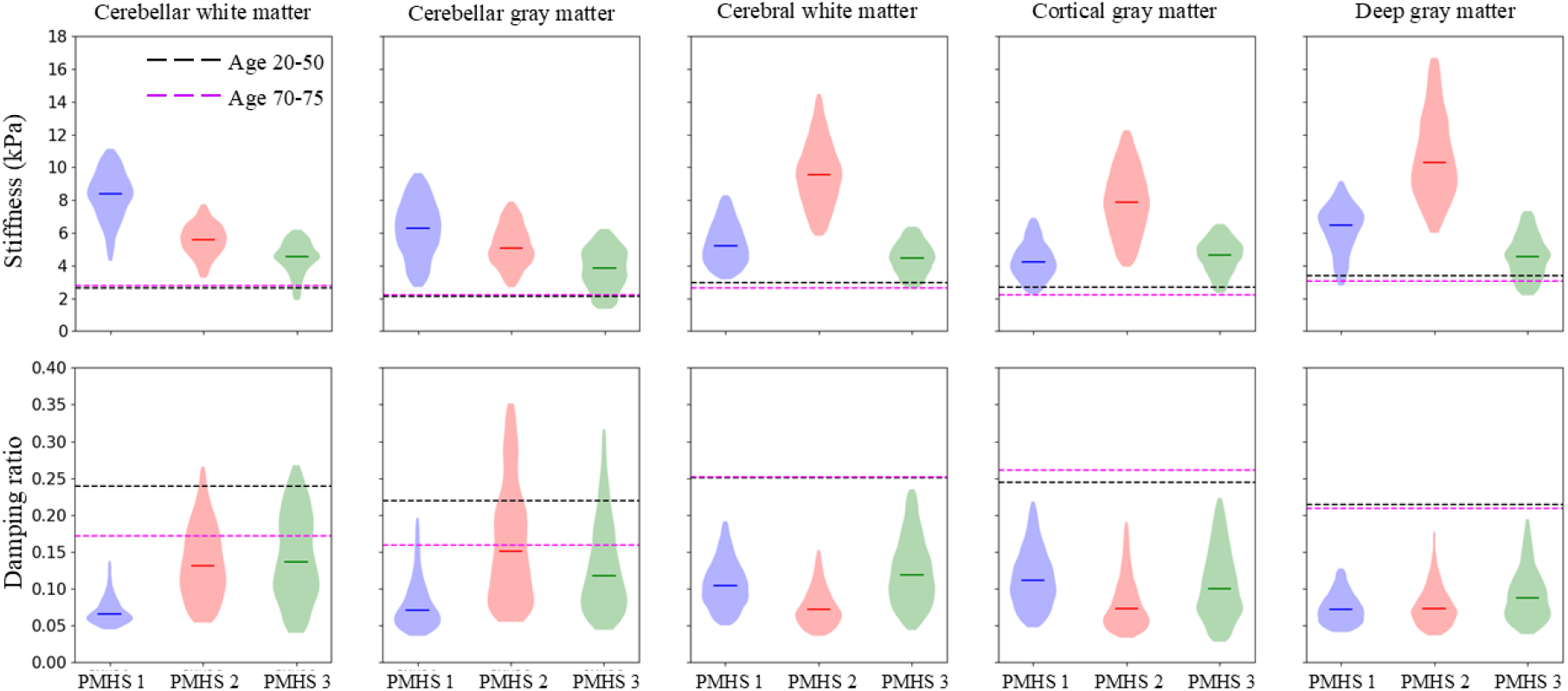
Violin plots of stiffness and damping ratio of cerebellar white matter, cerebellar gray matter, cerebral white matter, cortical gray matter, and deep gray matter. Median values of respective properties for *in vivo* subjects were shown with dotted lines. Data from corresponding regions for in vivo subjects are shown in **Figure A.1** in appendix.

Comparing the mechanical properties of the cerebrum and cerebellum, the white matter in the cerebellum was stiffer than white matter in cerebrum in PMHS 1 (61% stiffer) and PMHS 3 (2% stiffer) model, the white matter in the cerebellum was found to be 41% less stiff than white matter in cerebrum in both PMHS 2 model. On the other hand, the damping ratio of white matter was higher in cerebellar than cerebrum in PMHS 2 and PMHS 3 models, but lower in PMHS 1.

Regarding the gray matter properties, the stiffness was higher in deep gray matter followed by cerebellar gray matter and cortical gray matter in PMHS 1. However, in PMHS 2, deep gray matter was stiffest followed by cortical gray matter and then cerebellar gray matter. In PMHS 3, cortical gray matter was the stiffest followed by deep gray matter and then cerebellar gray matter. The damping ratio, on the other hand, was lowest in cerebellar gray matter and highest in cortical gray matter in PMHS 1. For PMHS 2 and 3, damping ratio was highest in cerebellar gray matter and lowest in deep gray matter. The quantitative results are tabulated in appendix A.1 and A.2.

### 3.3 Longitudinal post-mortem mechanical properties in PMHS 1

Figure 6 shows longitudinal changes in a representative axial slice of the stiffness and damping ratio, depicting non-uniform spatial variations and nonlinear trends. White matter and deep brain structures generally exhibited more changes over time than cortical regions. As shown in Table A.3, the brain tissue was stiffest on day 7 and softest on day 58. On the other hand, damping ratio was lowest on day 7 and highest on day 58.

**Figure 6.**
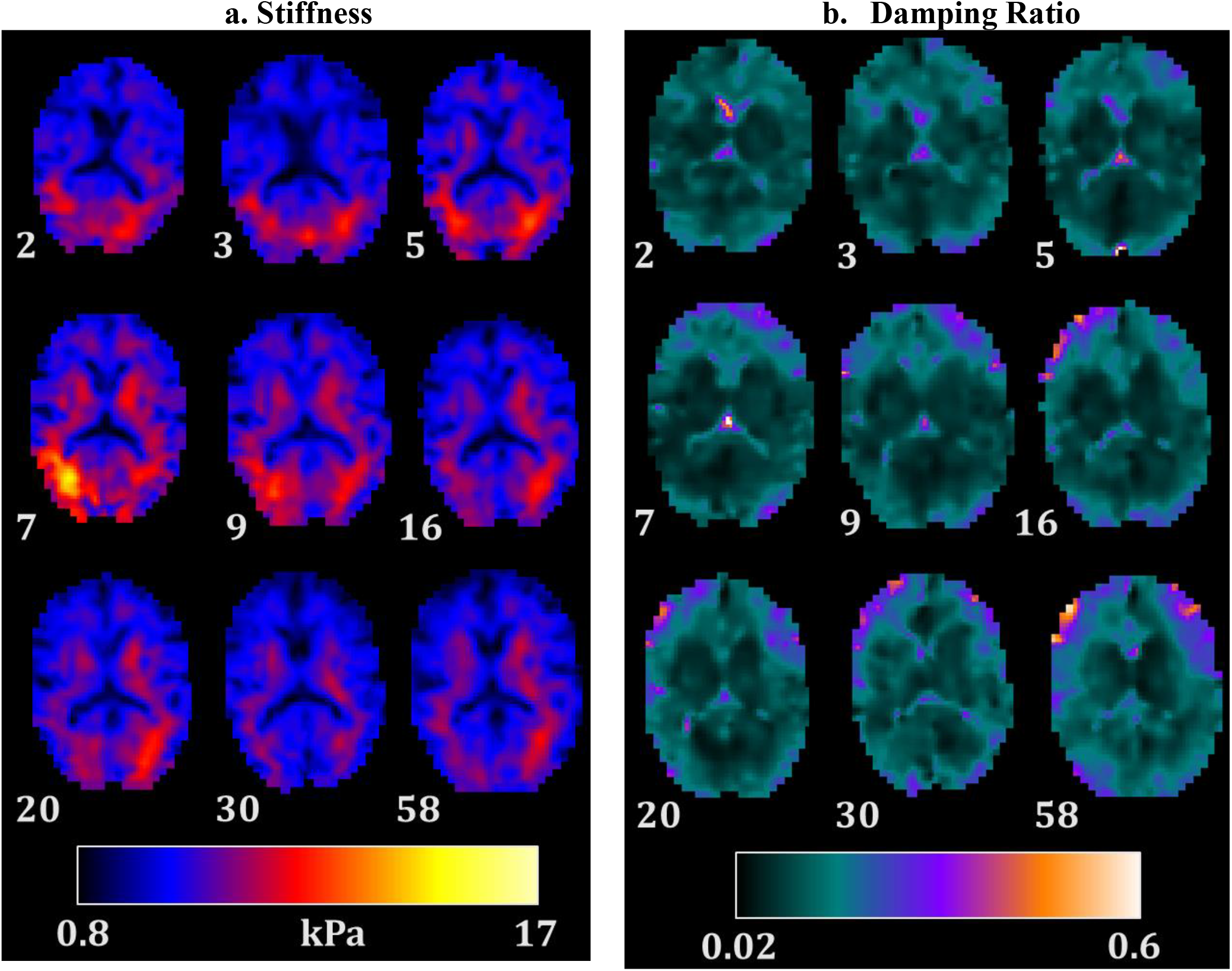
Spatial distribution of (**a)** stiffness and (**b)** damping ratio from day 2 to day 58. Values within the 5^th^ to 95^th^ percentiles are presented. The timepoint is shown to the bottom left of the image.

To illustrate the changes in properties of PMHS 1 over the post-mortem interval, Figure 7 shows the mechanical properties with all values normalized by the day 2 results. The figure shows that overall, the properties follow an apparent non-linear trend with inflections around days 3 and 7. Storage modulus, loss modulus, and stiffness increased on day 5 and day 7 compared to day 2 whereas gradual decreases in the same respective properties were found in days 9 to 58. On the other hand, the damping ratio decreased on days 5 and 7 compared to day 2, with gradual increases in from day 9 to day 58. Data on day 3 showed lower storage modulus, and stiffness, whereas damping ratio and loss modulus were higher than days 2 and 5.

**Figure 7.**
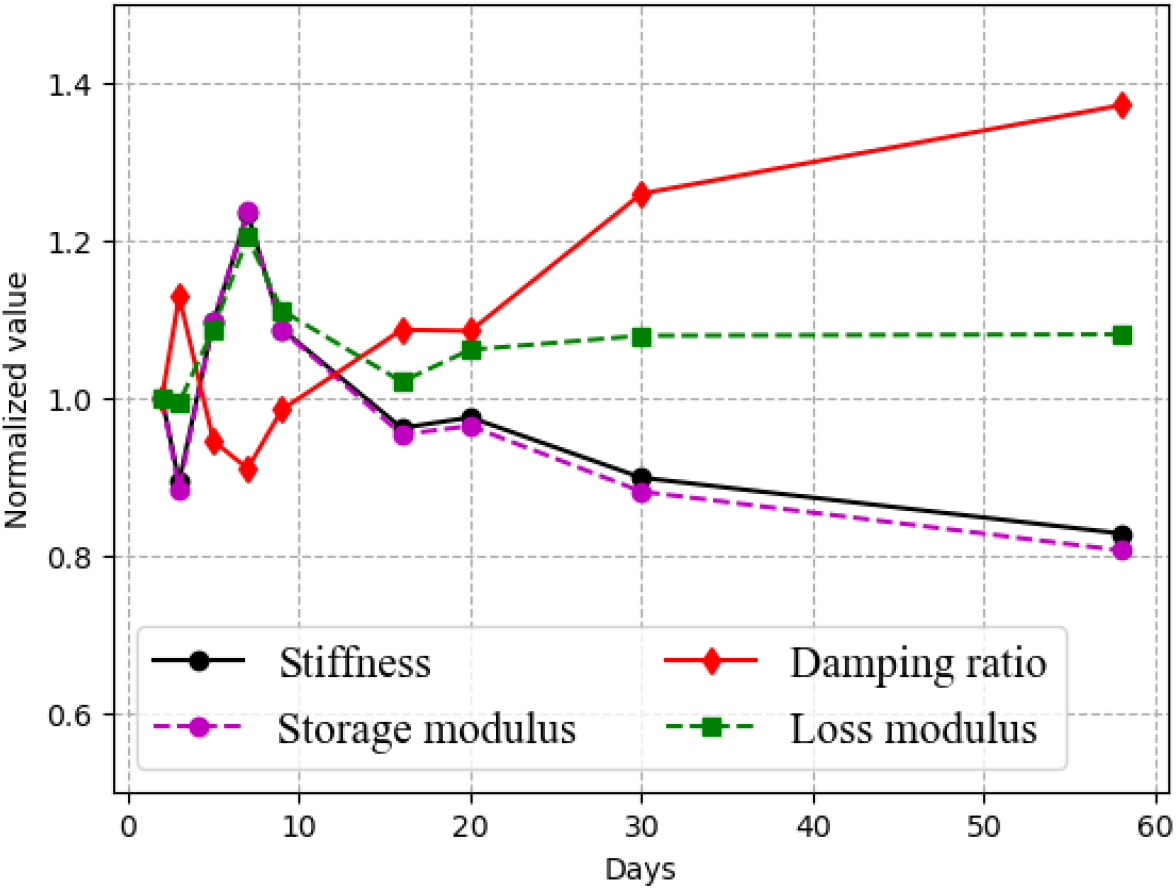
Median values of mechanical properties at respective days were normalized to the median values from day 2 for PMHS 1.

Figure 8 shows the longitudinal changes in regional stiffness and damping ratio of PMHS 1. Cerebral tissues were stiffest at day 7 (*μ* = 5.9 ± 1.34 *kPa*) whereas cerebellum was stiffest at day 5 (*μ* = 11.8 ± 4.9 *kPa*). In the cerebrum, white matter was slightly stiffer than cortical gray matter at day 2, but then the deep gray matter increased more rapidly than other regions through day 7. At day 58 cerebellar stiffness was at its lowest (*μ* = 4.61 ± 1.37*kPa*). A similar trend was observed in storage modulus. Damping ratio was lowest in the cerebellum (*ξ* = 0.03 ± 0.013) at day 5 and highest in the cerebrum (*ξ* = 0.14 ± 0.03) at day 58 (Figure 8). Cerebellar white matter was found to be stiffer than gray matter at all time points, whereas the relationship between white matter and gray matter in the cerebrum was more varied.

**Figure 8.**
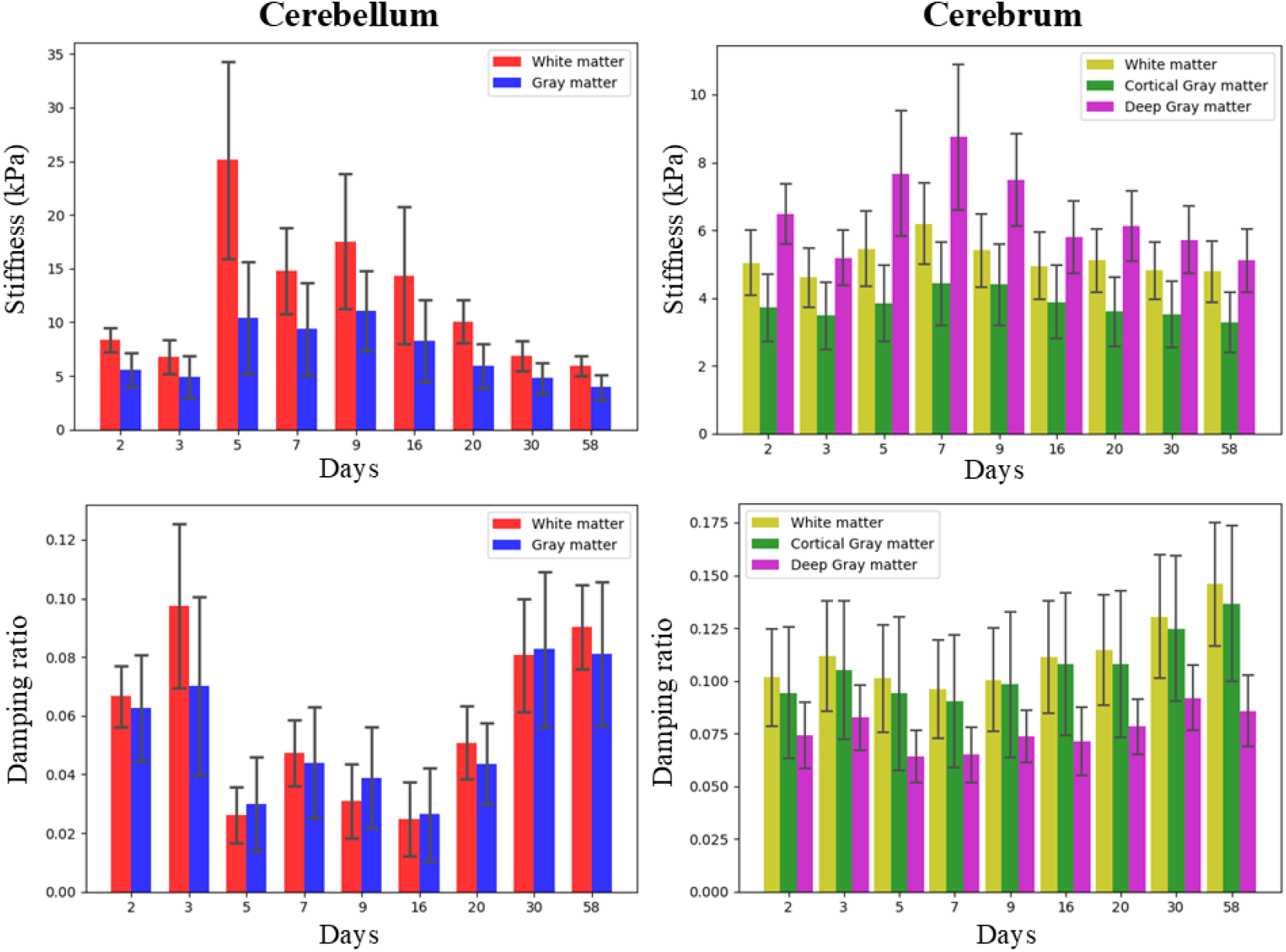
Bar plots showing the variation of material properties of white matter vs. gray matter in cerebellar (left column) and cerebrum (right column) from day 2 to day 58.

## 4 Discussion

In this study, we examined the post-mortem degradation of brain tissue mechanical properties by conducting MRE on three PMHS within 48-72 hours after death, one of which was scanned over an extended duration of nearly two months. The primary discovery of our investigation indicates an increase in stiffness and a decline in damping ratio within 48-72 hours post-mortem relative to *in vivo* subjects. Furthermore, our study suggests a nonlinear trend of tissue material properties over a time period of about two months. Specifically, we observed a decrease in stiffness and an increase in damping ratio in PMHS 1 after the seventh day post-mortem that was a reversal of the trends over the first seven days.

The initial increase in shear stiffness and a decrease in damping ratio in postmortem brain specimens indicates a shift towards greater elastic properties and reduced viscous properties. These findings are consistent with prior research conducted on porcine and mouse brain tissue (Gefen & Margulies, 2004; Miller et al., 2000), such as in one MRE study on eight minipig brains revealing a 145% increase in shear stiffness within an hour post-mortem compared to a live brain (Wang et al., 2024). Another MRE study on a single minipig brain reported an immediate brain stiffening up to 58% within three minutes post-mortem and a continued stiffening up to 142% within 45 mins (Weickenmeier et al., 2018). In addition, an MRE study on mouse brain reported an increase in stiffness after respiratory arrest and up to a 30% increase in cerebral stiffness within 5 mins of cardiac arrest (Bertalan et al., 2020). Our analysis also revealed some variability in material properties across all three models, suggesting possible subject-specific differences in mechanical properties. Notably, PMHS 2 exhibited a significant elevation in shear stiffness and a reduction in damping ratio compared to PMHS 1 and 3. This difference may be attributed to factors such as underlying tissue pathology prior to death, age-related changes in brain tissue, postmortem specimen handling, and the rate of tissue degradation following death. Additionally, in PMHS 1, we noted strong fluctuations in properties the first week after death, indicating that the time interval from death can substantially affect measurements.

This study also revealed an initial significant increase in many of the mechanical properties of the brain, including shear stiffness, storage modulus, and loss modulus, up to the seventh day post-mortem, followed by a decrease by the fifty-eighth day. These changes were not uniform across brain regions or between white matter and gray matter. Specifically, white matter in the cerebellum and cerebrum initially showed increased stiffness, which then declined, with the cerebellum experiencing a greater reduction. These trends are consistent with alterations in brain tissue mechanics over time after death found in rat brain tissue (Vappou et al., 2008). In that study, with the application of MRE on rat brain, Vappou et al found 100% increase in storage modulus right after death, whereas 50% decrease in loss modulus 24h after death. However, factors such as storage conditions, the method of euthanasia involving intracardiac injection of barbiturates that induces apnea and promotes blood influx into vital organs including the brain, as well as variations in brain size and inherent anatomical differences, may contribute to a more rapid alteration of tissue properties in the rat brain (Vappou et al., 2008).

The time elapsed between death and data acquisition should be considered a critical parameter in PMHS-based experimentation. Previous TBI studies (Giudice et al., 2019) aiming to elucidate brain injury mechanisms under injurious loading conditions have employed various techniques, such as biplanar X-ray imaging(Hardy et al., 2001; King et al., 2011) and sonomicrometry (Alshareef et al., 2020) on PMHS brains. While the sonomicrometry study reported testing within 72 hours post-mortem, the biplanar X-ray study did not specify the acquisition timeframe. Notably, even within a 72-hour postmortem window, our results demonstrated substantial variability in brain tissue mechanical properties across three PMHS models. Although these foundational experimental studies have been instrumental in validating computational TBI modes (Atsumi et al., 2018; Fernandes et al., 2018; Giudice et al., 2019), caution is warranted when extrapolating material property relationships between PMHS and in vivo human data, particularly in the context of injury biomechanics. Our results showed that PMHS 3 demonstrated shear stiffness and damping ratio closest to *in vivo* values. This specimen was the youngest of the three tested, indicating that it may be possible to optimize PMHS experiments for *in vivo* biofidelity based on demographics and time interval from death.

The mechanisms responsible for changes in the mechanical properties of the post-mortem brain are not yet fully understood. These alterations are likely associated with the depolarization of cell membranes that occurs shortly after death, leading to an influx of water and resulting in the swelling of neurons and astroglial cells (Risher et al., 2012; Santos et al., 2014; Thrane et al., 2014; Vappou et al., 2008). The observed reduction in brain tissue damping may stem from this cellular swelling, which reduces the volume of the extracellular space, and limits the mobility of extracellular water (Krassner et al., 2023). In addition, the loss of cerebral perfusion following death exerts both direct and indirect effects on brain mechanics under ischemic conditions. In living tissue, reduced perfusion is often linked to diminished stiffness. However, the complete cessation of cerebral blood flow after death may lead to cell damage or death, thereby disrupting cellular structure and compromising the integrity of the extracellular matrix (Doyle et al., 2008; Hetzer et al., 2018). Cerebellar changes were distinct from those in the cerebrum, perhaps highlighting the role of the different geometries and attachments, with the cerebellum being separated from the cerebrum by the tentorium cerebelli and connected to the spinal cord through the brainstem.

This study has several limitations. First, the exploration of mechanical properties was conducted using only three PMHS. Given the subject-specific nature of material degradation and property variation, future research could benefit from incorporating more PMHS to better understand the expected mechanical properties and their variability. Similarly, the study’s examination of changes in material properties over an extended period post-mortem was limited to a single PMHS. Despite the significant challenges and costs associated with obtaining and characterizing PMHS, future studies might consider shorter observation periods, such as up to a month after death, to enhance the applicability of the results. Although the earliest timepoints were greater than 24 hours after death, this is likely the amount of time that would be required in most experimental PMHS studies (c.f. Table 1 from Alshareef et al (Alshareef et al., 2020)). Secondly, material properties were estimated using the NLI method, based on the assumption that the brain behaves as an isotropic, hyper-elastic material. Notably, artifactual hot spots may occur due to excessive phase wraps in MRE. Advancements in the estimation method could accommodate anisotropic materials, potentially leveraging diffusion tensor imaging to ascertain directionality (Basser & Pierpaoli, 1996; Feng et al., 2013; Tweten et al., 2017). Lastly, further research is needed to uncover the cellular and biological mechanisms driving material degradation post-mortem.

## 5 Conclusion

This study marks the first detailed investigation into the post-mortem changes in the mechanical properties of human brain tissue. It assessed the mechanical properties derived from wave displacement fields at various post-mortem intervals, comparing them with those of living human subjects, indicating an initial increase in brain tissue stiffness, storage modulus, and wave speed, alongside a decrease in damping ratio and viscosity. Over time, stiffness initially increased then decreased, while the storage modulus and wave propagation speed followed a similar pattern, contrasting with the damping ratio and viscosity trends. These findings could contribute to a better understanding of the limitations of using PMHS in biomechanics experiments, which are important for studies involving traumatic brain injury and neurosurgery.

## 6 Acknowledgements

This work was partially supported by NINDS grant R01-NS136056, NINDS grant U01-NS112120, the Department of Defense in the Military Traumatic Brain Injury Initiative (MTBI2), the intramural research program of the NIH and the NIH Bench-to-Bedside and Back Program. We greatly appreciate Dr. Andrew Knutsen’s insights on this study. The authors have no conflicts of interest to disclose. The opinions and assertions expressed herein are those of the author(s) and do not reflect the official policy or position of the Henry M. Jackson Foundation for the Advancement of Military Medicine, Inc., the National Institutes of Health, the Uniformed Services University of the Health Sciences, or the Department of Defense.

## 7 Appendix

**Table A.1.**
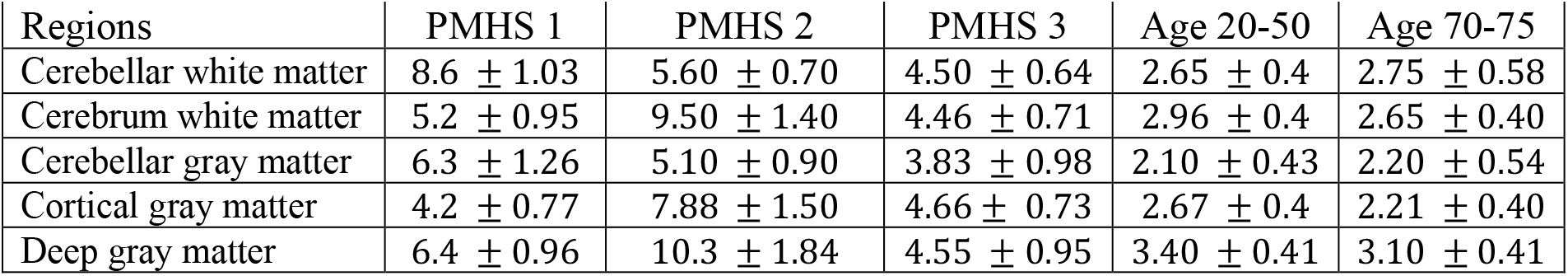
Regional stiffness (kPa) (median ± median absolute deviation) in PMHS models.

| Regions | PMHS 1 | PMHS 2 | PMHS 3 | Age 20-50 | Age 70-75 |
| --- | --- | --- | --- | --- | --- |
| Cerebellar white matter | 8.6 ± 1.03 | 5.60 ± 0.70 | 4.50 ± 0.64 | 2.65 ± 0.4 | 2.75 ± 0.58 |
| Cerebrum white matter | 5.2 ± 0.95 | 9.50 ± 1.40 | 4.46 ± 0.71 | 2.96 ± 0.4 | 2.65 ± 0.40 |
| Cerebellar gray matter | 6.3 ± 1.26 | 5.10 ± 0.90 | 3.83 ± 0.98 | 2.10 ± 0.43 | 2.20 ± 0.54 |
| Cortical gray matter | 4.2 ± 0.77 | 7.88 ± 1.50 | 4.66 ± 0.73 | 2.67 ± 0.4 | 2.21 ± 0.40 |
| Deep gray matter | 6.4 ± 0.96 | 10.3 ± 1.84 | 4.55 ± 0.95 | 3.40 ± 0.41 | 3.10 ± 0.41 |

**Table A.2.** Regional damping ratio (median ± median absolute deviation) in PMHS models.

| Regions | PMHS 1 | PMHS 2 | PMHS 3 | Age 20-50 | Age 70-75 |
| --- | --- | --- | --- | --- | --- |
| Cerebellar white matter | 0.07 ± 0.01 | 0.13 ± 0.04 | 0.14 ± 0.05 | 0.24 ± 0.05 | 0.17 ± 0.04 |
| Cerebrum white matter | 0.10 ± 0.02 | 0.07 ± 0.02 | 0.12 ± 0.03 | 0.25 ± 0.05 | 0.25 ± 0.05 |
| Cerebellar gray matter | 0.07 ± 0.02 | 0.15 ± 0.06 | 0.12 ± 0.04 | 0.22 ± 0.06 | 0.16 ± 0.05 |
| Cortical gray matter | 0.10 ± 0.03 | 0.07 ± 0.02 | 0.10 ± 0.03 | 0.24 ± 0.06 | 0.26 ± 0.07 |
| Deep gray matter | 0.07 ± 0.02 | 0.07 ± 0.02 | 0.09 ± 0.02 | 0.21 ± 0.04 | 0.21 ± 0.04 |

**Table A.3.** Mechanical properties of whole brain in PMHS 1 over time.

| Index | Days | Stiffness (kPa) | Damping ratio |
| --- | --- | --- | --- |
| 1 | 2 | 4.80 ± 1.20 | 0.09 ± 0.03 |
| 2 | 3 | 4.30 ± 1.10 | 0.11 ± 0.03 |
| 3 | 5 | 5.30 ± 1.50 | 0.09 ± 0.03 |
| 4 | 7 | 5.94 ± 1.66 | 0.08 ± 0.03 |
| 5 | 9 | 5.23 ± 1.43 | 0.09 ± 0.03 |
| 6 | 16 | 4.63 ± 1.24 | 0.10 ± 0.03 |
| 7 | 20 | 4.70 ± 1.26 | 0.10 ± 0.03 |
| 8 | 30 | 4.30 ± 1.14 | 0.12 ± 0.03 |
| 9 | 58 | 3.98 ± 1.13 | 0.13 ± 0.04 |

---

**Figure A.1.**
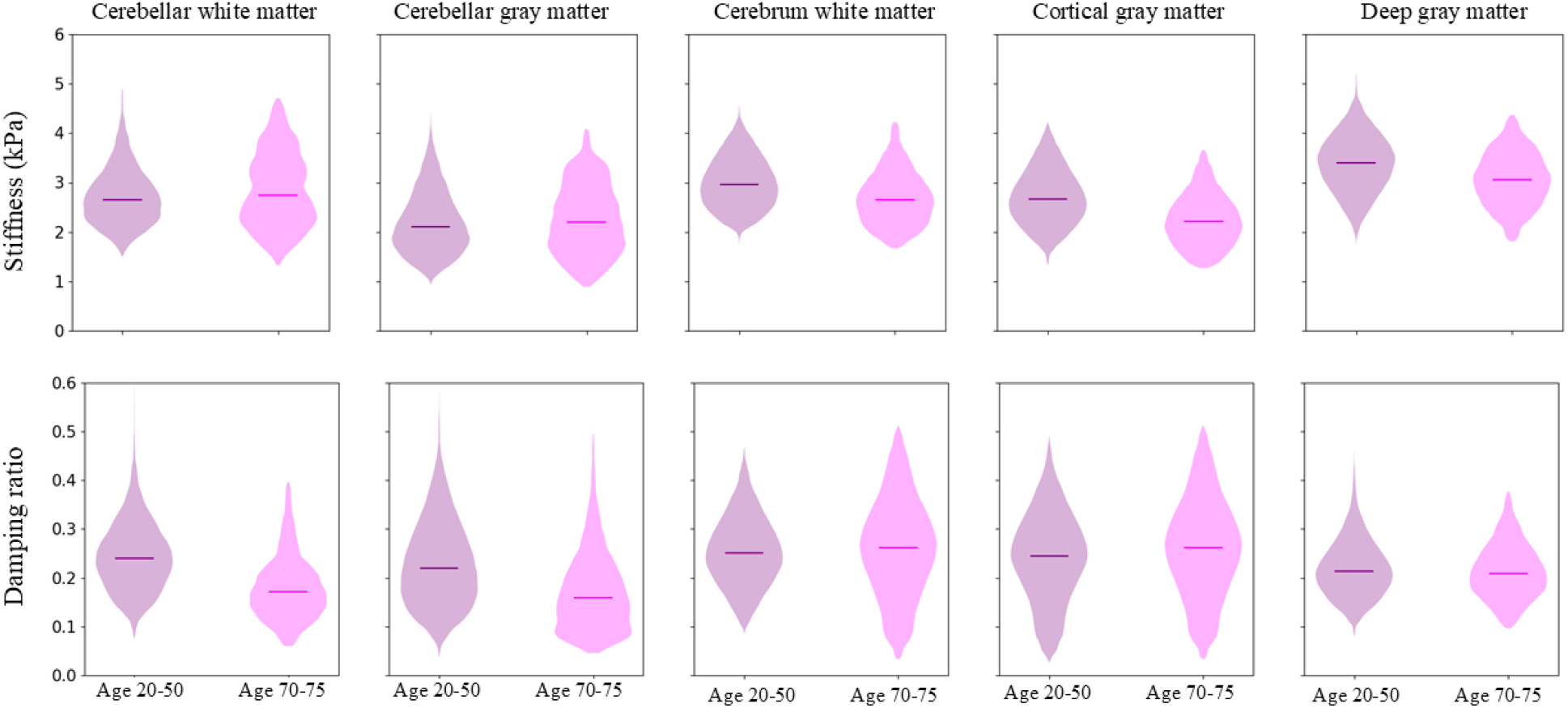
Violin plots of stiffness and damping ratio of cerebellar white matter, cerebellar gray matter, cerebral white matter, cortical gray matter, and deep gray matter in *in-vivo* subjects.

